# A novel *in silico* method employs chemical and protein similarity algorithms to accurately identify chemical transformations in the human gut microbiome

**DOI:** 10.1101/2022.08.02.502504

**Authors:** Annamarie Bustion, Ayushi Agrawal, Peter J. Turnbaugh, Katherine S. Pollard

**Author notes:** Correspondence to: Katherine Pollard, 1650 Owens Street, San Francisco, CA 94158.

## Abstract

Bacteria within the gut microbiota possess the ability to metabolize a wide array of human drugs, foods, and toxins, but the responsible enzymes for these chemical events remain largely uncharacterized due to the time consuming nature of current experimental approaches. Attempts have been made in the past to computationally predict which bacterial species and enzymes are responsible for chemical transformations in the gut environment, but with low accuracy due to minimal chemical representation and sequence similarity search schemes. Here, we present an *in silico* approach that employs chemical and protein <u>S</u>imilarity algorithms that <u>I</u>dentify <u>M</u>icrobio<u>M</u>e <u>E</u>nzymatic <u>R</u>eactions (SIMMER). We show that SIMMER predicts the chemistry and responsible species and enzymes for a queried reaction with high accuracy, unlike previous methods. We demonstrate SIMMER use cases in the context of drug metabolism by predicting previously uncharacterized enzymes for 88 drug transformations known to occur in the human gut. Bacterial species containing these enzymes are enriched within human donor stool samples that metabolize the query compound. After demonstrating its utility and accuracy, we chose to make SIMMER available as both a command-line and web tool, with flexible input and output options for determining chemical transformations within the human gut. We present SIMMER as a computational addition to the microbiome researcher’s toolbox, enabling them to make informed hypotheses before embarking on the lengthy laboratory experiments required to characterize novel bacterial enzymes that can alter human ingested compounds.

## Introduction

Humans consume a large array of foods, therapeutics, and other xenobiotics that are processed, in part, by enzymes of bacteria residing within the gut. While some bacterial enzymes are orthologous to the human metabolism repertoire, many bacteria possess metabolic capabilities distinct from our own (Zimmermann et al., 2019a). It is important to ascertain the extent of microbial capacity for chemical transformation because it has implications for the bioavailability, toxicity, and efficacy of the compounds humans ingest (Koppel et al., 2017; Spanogiannopoulos et al., 2016). Additionally, because the human gut microbiome differs from individual to individual, knowledge of the prevalence and abundance of bacterial enzymes must be determined before beneficial clinical and dietary decisions can be made (Javdan et al., 2020).

While experimental methods can be employed to expand what we know of bacterial enzymatic capabilities in the gut, the scientific community lacks genetic tools for nearly all bacterial species of the human microbiota, and heterologous expression in model organisms can fail for a plethora of reasons (Bisanz et al., 2020; Patel et al., 2022). When experimental methods are tractable, the time required is often so extended that knowledge is gained in a low-throughput manner. For these reasons, attention should turn to the employment of *in silico* computational methods that can guide experimentalists in their hypothesis-building process by aiding in the prioritization of substrates, species, and genes worth studying.

Recent attempts have been made to create computational descriptions of chemical transformation by human gut bacteria, but none can be expanded to predict the metabolic capabilities of bacterial proteins with unknown function or to explore the capacity of microbial enzymes to degrade novel substrates. Two previously published methods aimed to predict known drug metabolism events within the human gut microbiome, but the accuracy of their predictions was limited due to the fact that both tools only consider substrates, rather than a full chemical description of substrate(s), cofactor(s), and product(s) formed in a reaction. Both tools were also limited by the use of small databases that do not fully capture the diversity of the human gut microbiome (Guthrie et al., 2019; Sharma et al., 2017).

To address these gaps in accurate predictive software for bacterial chemical transformations, we present SIMMER, a tool that combines chemoinformatics and metagenomics approaches to accurately predict bacterial enzymes capable of metabolism events. Given an input reaction, SIMMER predicts an Enzyme Commision (EC) code that describes the chemical nature of the query. SIMMER additionally predicts specific bacterial enzyme sequences, functions, prevalence, and abundance for the reaction. Our key innovations are the use of full chemical representations that include cofactors employed and products produced in a reaction, the use of statistically informed sequence searches of a comprehensive human gut microbiome gene catalog, and the development of a novel EC predictor based on reaction rather than gene sequence. As a use-case, we present evidence that SIMMER provides high accuracy predictions of bacterial enzymes responsible for known drug metabolism events, and we identify the likely bacterial enzyme for 88 drugs known to be metabolized by the gut microbiome for which the enzyme was previously unknown.

## Results

### SIMMER pipeline to predict xenobiotic metabolizing enzymes

There are many desiderata for a bacterial drug metabolism predictor (Table 1). Such a tool must be able to, based on quantified chemical similarity, predict EC annotations, specific enzyme sequences, and the prevalence and abundance across human samples of those predicted sequences. We developed SIMMER, a tool that leverages chemical and protein similarity to identify enzymes in the human microbiome that could perform a queried chemical reaction (Figure 1). Given input substrate(s), metabolite(s), and any known cofactors, SIMMER predicts bacterial enzymes capable of performing the reaction and quantifies their prevalence and abundance in the human gut. SIMMER accomplishes this by chemically fingerprinting an input reaction, and then comparing it to all reactions in MetaCyc. Enzyme annotations from the most similar MetaCyc reactions are then used as queries for a protein similarity search to find homologs in the genomes of gut bacteria. To decrease the runtime of a SIMMER query, we precomputed chemical descriptions and protein similarity searches for all reactions in MetaCyc.

**Table 1.** Desired elements of a microbiome chemical transformation predictor.

|  |  | DrugBug<br>(Sharma et al.,<br>2017) | MicrobeFDT<br>(Guthrie et al.,<br>2019) | **SIMMER** |
| --- | --- | --- | --- | --- |
| Input types | Accepts novel SMILES |  |  | ✓ |
|  | User options for different chemical fingerprints | ✓ |  | ✓ |
| Output types | Reaction similarity measure |  |  | ✓ |
|  | EC predictions | ✓ | ✓ | ✓ |
|  | Enzyme and species predictions | ✓ |  | ✓ |
|  | Function predictions |  |  | ✓ |
|  | Prevalence/abundance |  | ✓ | ✓ |
| Usability | Web server | ✓ |  | ✓ |
|  | Command line tool |  |  | ✓ |
|  | Docker container |  | ✓ |  |
| Accuracy | Prediction of previously characterized enzymes | 3% | NA | **87%** |
|  | Prediction of previously characterized EC numbers | 39% | 14% | **93%** |

**Figure 1.**
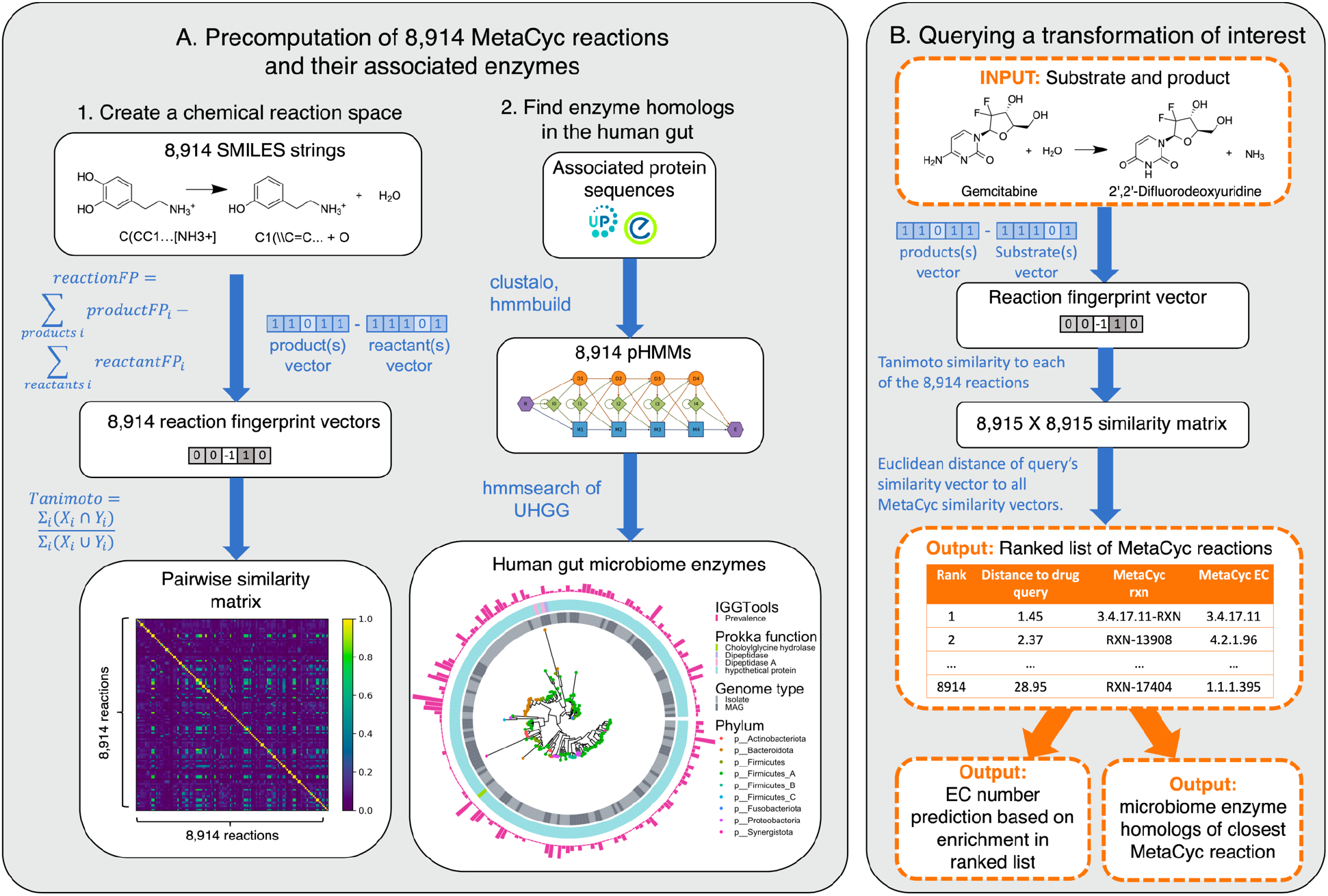
SIMMER architecture. **(A)** Precomputation on 8,914 gene annotated bacterial reactions downloaded from MetaCyc. Chemical fingerprints representing each MetaCyc reaction were created from SMILES descriptors. A latent chemical space was then created via a pairwise reaction similarity matrix based on Tanimoto coefficients. For each reaction, relevant gene sequences were retrieved from UniProt and Entrez database linkouts and used to create multiple sequence alignments and subsequent pHMMs using ClustalO and HMMER3 respectively. pHMMs were used to retrieve homologs in a catalog of human gut microbiome genes. **(B)** Running a SIMMER query. After receiving a reaction query (input compound, co-factors, products), SIMMER fingerprints the reaction and compares it to the precomputed chemical space from 1A. MetaCyc reactions are sorted by similarity to the query. Significantly enriched EC identities are reported, and from the most similar reaction, human gut microbiome enzymes are reported along with their abundance and prevalence in gut microbiomes.

SIMMER’s underlying data was drawn from the MetaCyc reaction database because its small-molecule reaction descriptions each possess at least one experimentally validated enzyme annotation and often include a description of the reaction type via EC code. To build a precomputed chemical search space for SIMMER queries (Figure 1A), we created two-dimensional fingerprint representations for 8,914 enzyme driven reactions in MetaCyc (Caspi et al., 2008; Schneider et al., 2015). Using these fingerprints, we estimated the similarity between all pairs of reactions based on Tanimoto coefficients. To build the enzyme backbone of SIMMER, we compiled the Uniprot and/or Entrez gene identifiers linked to each MetaCyc transformation into a profile hidden Markov model (pHMM) that represents the diversity of the enzyme family for a respective reaction. The resulting pHMMs were then used to query the Unified Human Gastrointestinal Genome (UHGG) collection of 286,997 isolate genomes and metagenome assembled genomes from the human gut environment (Almeida et al., 2020). Additionally, prevalence and abundance of all pHMM search hits were assessed in stool metagenomes from the PREDICT human cohort using MIDAS2, an implementation of Metagenomic Intra-Species Diversity Analysis System (MIDAS) designed for use with the UHGG catalog (Almeida et al., 2020; Nayfach et al., 2016; Zhao et al., 2022).

After creating SIMMER’s precomputed chemical and pHMM search space, we next made SIMMER queryable (Figure 1B). When queried with a chemical transformation, SIMMER computes the chemical similarity of the input to all precomputed MetaCyc reactions, and sorts all MetaCyc reaction fingerprints according to their ascending Euclidean distance from the query. From this sorted list, SIMMER outputs enzymes (i.e. the precomputed pHMM search hits for the closest reactions) responsible for the query reaction and an EC code (i.e. reaction type) prediction. We implemented and validated a novel method to predict EC codes by extending a common approach to gene set enrichment analysis (GSEA) (Figure 2—figure supplement 1) (Subramanian et al., 2005). With this enrichment method, SIMMER predicted reaction types for queries with high recall, precision, and accuracy for EC classes, sub-classes, and sub-sub-classes (Figure 2B, Figure 2—source data, Figure 2—figure supplement 1).

**Figure 2.**
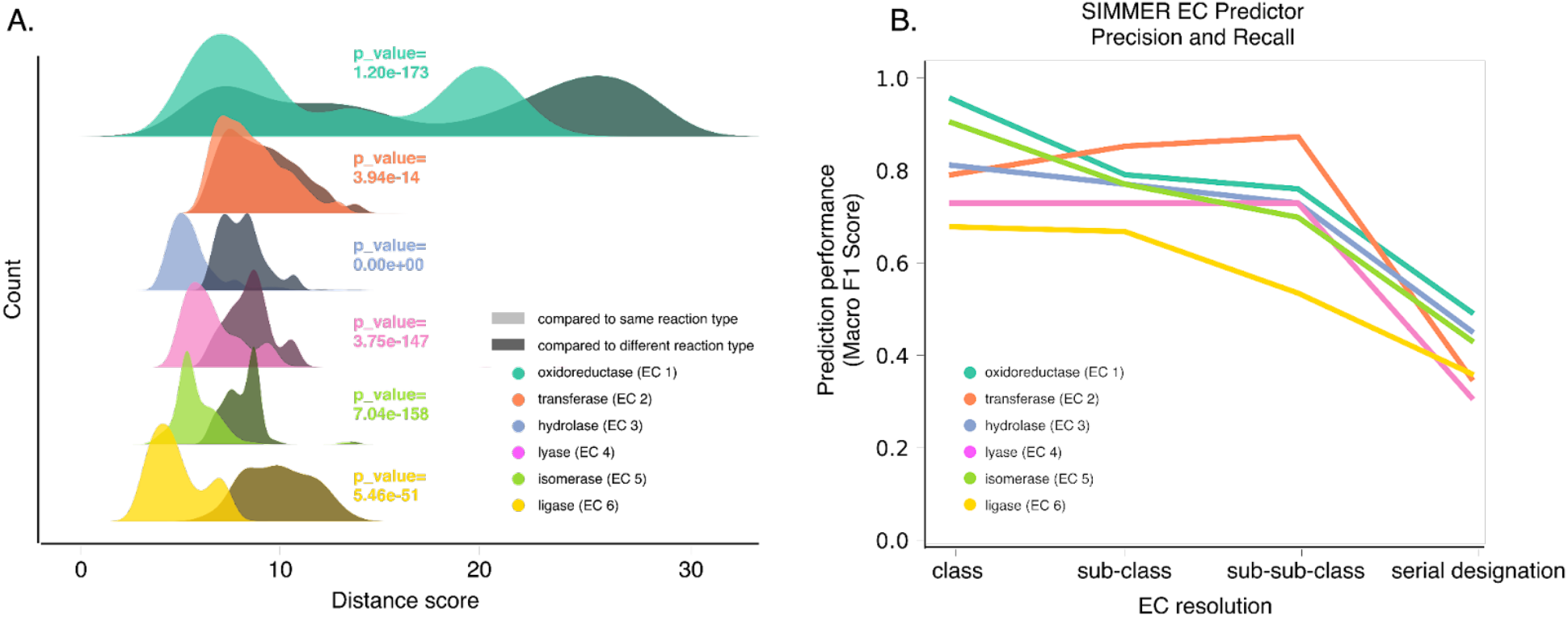
SIMMER’s chemical representations capture information relevant to enzymatic reactions. **(A)** SIMMER clusters similar reactions together in chemical space. To analyze SIMMER’s ability to group chemically similar reactions, we examined reaction similarity within versus between EC classes using the procomputed MetaCyc reaction dataset (N=8,914 reactions). A silhouette-like euclidean distance score was created by determining for each reaction its euclidean distance to all reactions within its EC class versus outside its EC class. For all EC classes, scores were smaller within versus between EC classes using SIMMER’s chemical representation, indicating that SIMMER can detect reaction similarity within EC classes. From the pairs of distributions we computed a Kolmogorov-Statistic to determine if the distributions significantly (p<0.05) differed. **(B)** The F1-score, or harmonic mean of SIMMER’s precision and recall, when predicting EC numbers on a subset of the MetaCyc database (N=576 reactions total; 96 per EC class). The score is high for EC classes, and it generally decreases as an EC number’s resolution increases.

We hypothesized that SIMMER accurately predicted chemical transformations due to its use of a full reaction that includes reactants, cofactors, and products, rather than just substrates. We assessed this hypothesis by demonstrating that SIMMER groups similar reactions together in chemical space. MetaCyc reactions possess EC annotations that describe the chemical class of a reaction (e.g. oxidoreduction, hydrolysis, intramolecular rearrangement, etc). We queried SIMMER with all EC annotated MetaCyc reactions and demonstrated that queries group significantly closer to other reactions within their EC class than they do to reactions of a different class (Figure 2A). We determined that SIMMER’s ability to group similar reactions in chemical space is resilient to different fingerprinting methods (Figure 2—figure supplement 2), but not to loss of products created and cofactors consumed in a reaction (Figure 2—figure supplement 3). Thus we showed that similar reactions only cluster together in chemical space when a full reaction description (i.e. SIMMER’s representation method) is employed.

Because SIMMER was created with the assumption that chemically similar reactions are mediated by sequence similar enzymes, we next ensured that similarity within SIMMER’s chemical space could be used to find shared, responsible enzymes. First, for all MetaCyc enzymes associated with multiple reactions, one reaction was used as a SIMMER query, and the second reaction searched for in the ordered reaction list output. As a negative control, these reaction similarity results were then compared to all possible pairwise combinations of reactions not conducted by the same enzyme. SIMMER predicted high similarity between reactions conducted by a shared enzyme, and low similarity for those reactions without a shared enzyme, (Figure 3A). We also found a negative association between chemical reaction distance and sequence similarity of MetaCyc enzyme annotations, indicating that reactions with similar chemistry are conducted by sequence similar enzymes, though there is much variation in this relationship (Figure 3B). This reflects the known association between sequence similarity and similarity in chemical function, as well as reports that this relationship can often be overestimated (Tian and Skolnick, 2003). Together, these analyses demonstrated that sequence similar enzymes do indeed mediate chemically similar reactions, strengthening the logic of combining chemical and protein similarity in a microbiome enzyme prediction tool.

**Figure 3.**
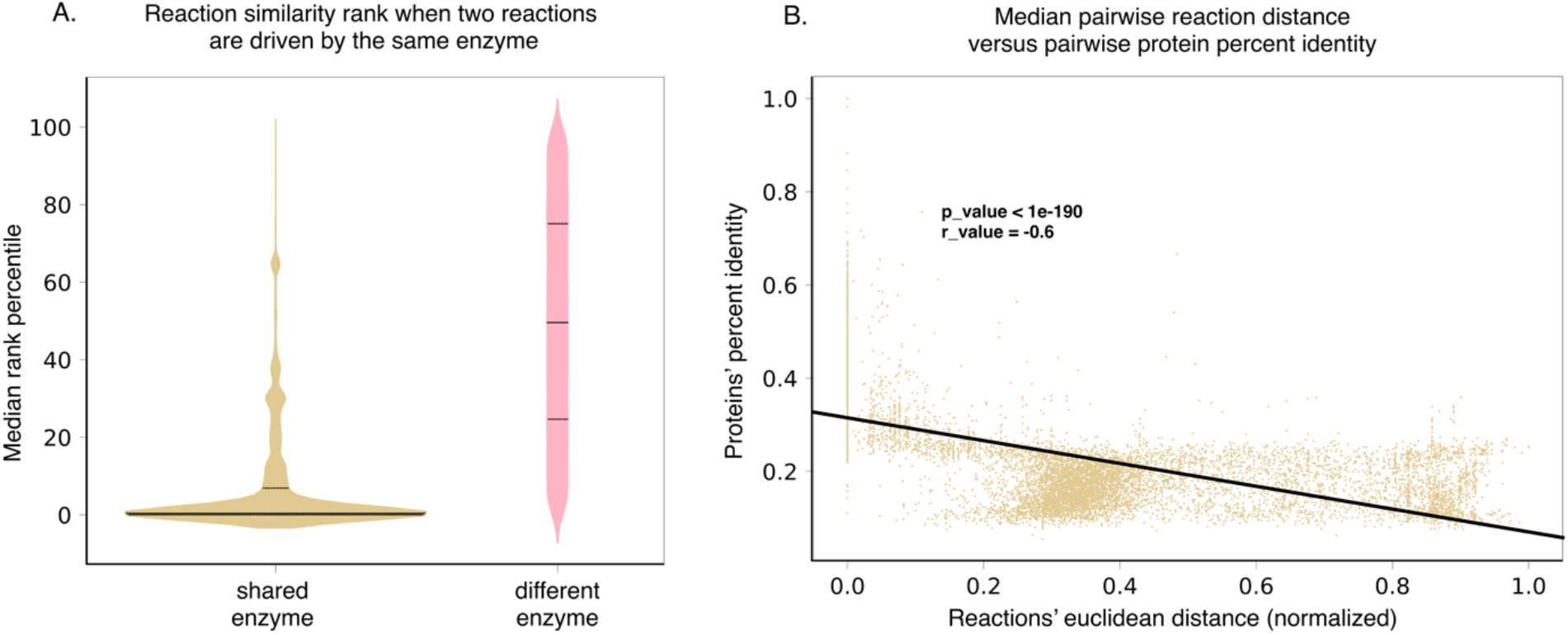
SIMMER’s chemical representations can be used to find shared, responsible enzymes. **(A)** When SIMMER was queried with a MetaCyc reaction, other reactions driven by the same enzyme are returned as the most similar. As a contrast, reactions driven by a different enzyme yield a more uniform rank distribution. Solid lines of the violin plots depict median reaction similarity rank and dashed lines represent lower and upper quartile ranges. **(B)** Similarly, there is a negative association between pairwise reaction euclidean distance and pairwise protein identity, demonstrating that SIMMER can capture the known, albeit weak relationship between sequence identity and similar reaction chemistry (Tian and Skolnick, 2003).

### An expanded list of gut bacterial enzymes relevant to known cases of drug metabolism

To assess SIMMER’s prediction accuracy for previously characterized reactions and to mount a comparison to existing methods, we used drug metabolism as a use case. First, we curated 298 drug-metabolism events associated with the human gut microbiome from the literature (Supplementary File 1). For 31 of these reactions the responsible bacterial enzyme, characterized metabolite(s), and associated EC annotation are known (Supplementary File 1). These 31 reactions are conducted by 18 enzymes. Due to orthology and proclivity for genetic transfer between even distantly related bacteria, however, there are likely many as yet undiscovered homologs of these drug-metabolizing enzymes that can catalyze identical drug metabolism events (Pollet et al., 2017). To account for this, we created an expanded database (Figure 4, Figure 4—figure supplement 1, Figure 4—source data) of the 18 characterized enzymes from pHMM and phmmer searches of the UHGG database (Almeida et al., 2020), yielding 52,849 total candidate homologs (a median of 1,087 candidates per enzyme). After filtering enzymes by hmmer significance, alignment length, presence in data from the human jejunum (Zmora et al., 2018) and RNA-sequencing studies (Integrative HMP (iHMP) Research Network Consortium, 2019), and predicted affinity for the substrate in question using the Similarity Ensemble Approach (Keiser et al., 2007), our database contained a median of 2 high-confidence homologous sequences per enzyme (range = 0 to 460 across the 18 enzyme families, Figure 4, Figure 4—figure supplement 1, Figure 4— source data). These 741 additional enzyme sequences for 31 reactions formed our positive control set of known gut bacterial enzymes.

**Figure 4.**
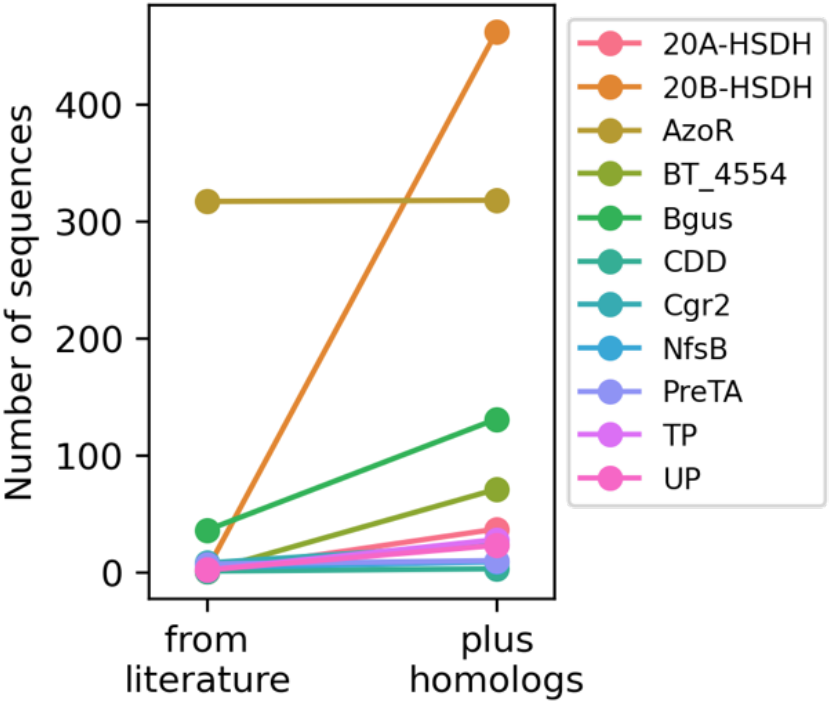
An expanded list of gut bacterial enzymes relevant to known cases of drug metabolism. Eleven of the 18 enzymes responsible for positive control drug-metabolism events have high confidence homologs that we gathered by filtering for biological significance.

### SIMMER captures known gut bacterial enzymes involved in drug metabolism

With our expanded database of drug-metabolizing enzymes from the human gut microbiome in hand, we next verified that SIMMER can accurately predict reaction types and responsible enzymes for the 31 known chemical transformations. Only 3 of these 31 reactions are themselves MetaCyc entries (5-ASA, dopamine, and levodopa degradation); if EC codes and enzymes of reactions not described in MetaCyc were also accurately predicted, it would show that SIMMER can discover non-identical yet chemically similar reactions.

Of the 31 drug-metabolism events known to occur via human gut bacterial enzymes, EC annotations exist for 28. SIMMER identified the correct EC class for 26 of these 28 reactions (93%) (Figure 5, Figure 5—source data). For some queries SIMMER predicted more than one significant EC code, but again, for 26 out of the 28 reactions, the top EC class prediction was a match (Figure 5, Figure 5—figure supplement 1, Figure 5—source data). The two failed EC predictions were for nicardipine reduction (inappropriately predicted as an isomerase reaction) and for brivudine transformation (for which SIMMER made no significant prediction).

**Figure 5.**
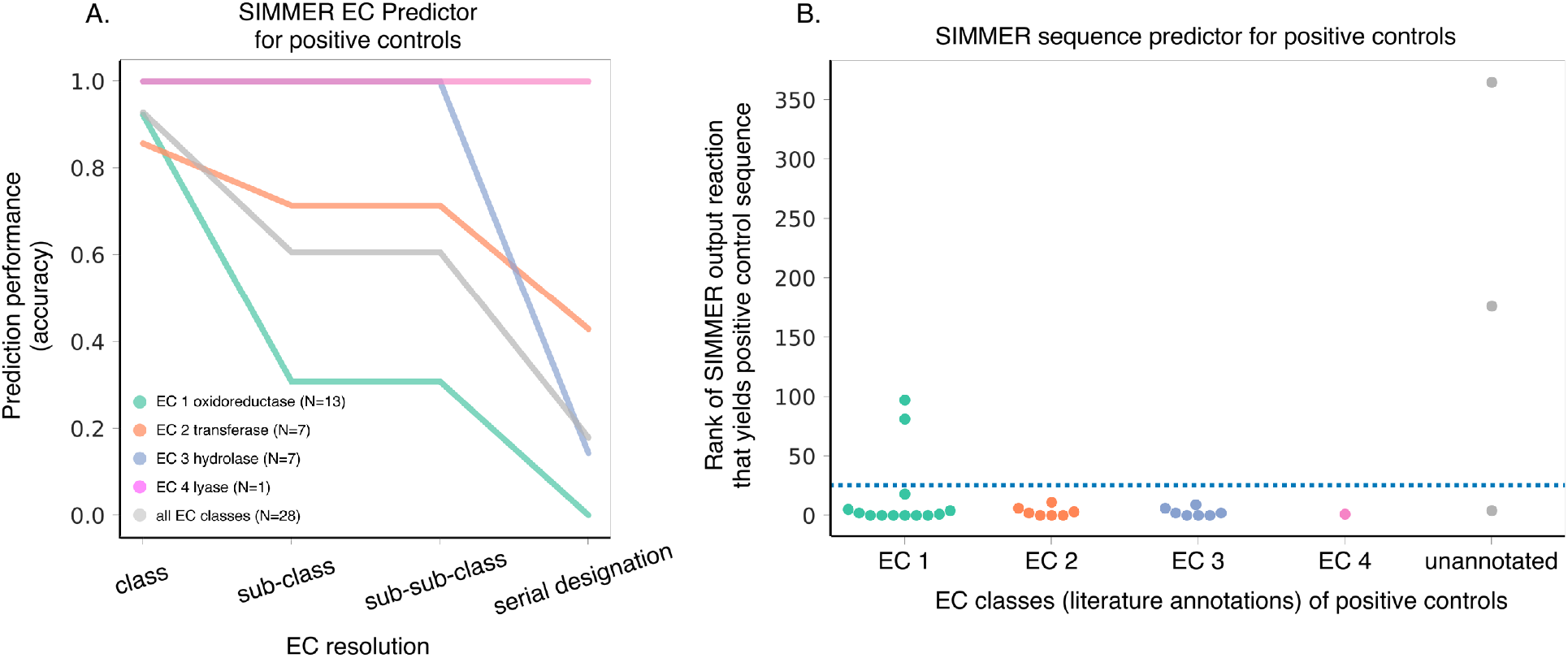
SIMMER captures known gut bacterial enzymes involved in drug metabolism. **(A)** SIMMER accurately predicted EC classes for 28 previously characterized reactions that possess EC annotations. As with the MetaCyc database (Figure 2B), accuracy dropped off as EC resolution increased. **(B)** SIMMER predicted bacterial sequences previously shown to drive 31 drug-metabolism events in the gut microbiome. Depicted is the rank (out of N=8,914 reactions) of the MetaCyc reaction that yielded a gut microbiome homolog matching the known positive control sequence. Reported accuracy is based on such a reaction being within the top 25 ranked reactions (dashed blue line).

In addition to accurate EC (i.e. reaction type) identification, SIMMER also accurately predicted the specific enzymes from the human gut microbiome that conduct the 31 query reactions (Supplementary File 4). This enzyme list was populated by the results of the precomputed pHMM searches of human microbiome catalogs with annotated gene sequences from MetaCyc reactions (Figure 1A). In 27 cases (87%), the characterized (i.e. positive control) enzyme(s) for a reaction was found in the output enzyme list for the top 20 of the ranked MetaCyc reactions (Figure 5—figure supplement 1). Since the positive controls span four EC classes (EC1 oxidoreductases, EC2 transferases, EC3 hydrolases, and EC4 lyases) this result demonstrates SIMMER’s ability to accurately predict microbiome based enzymes for a diversity of reaction types. Also, despite inaccurate EC predictions for nicardipine reduction and brivudine transformation, SIMMER was able to respectively predict AzoR and BT_4554 enzymes as responsible for the reactions (Figure 5, Figure 5—figure supplement 1, Figure 5—source data).

### SIMMER outperforms existing methods

To mount a comparison to the other *in silico* methods that, in part, aimed to describe microbiome drug metabolism, we next queried the 31 positive control reactions using MicrobeFDT and DrugBug (Table 1) both of which rely solely on substrate chemical similarity rather than information from a whole reaction (Guthrie et al., 2019; Sharma et al., 2017). For the 28 EC annotated positive control reactions, DrugBug had 39% accuracy in predicting EC classes (in comparison to SIMMER’s 93%), and predicted the correct enzyme for a single reaction, SN38 glucuronide deconjugation, despite the presence of chemically similar reactions metabolized by the same enzyme amongst the positive controls (Table 1—source data). We additionally queried SIMMER with the four positive controls (ginsenoside Rb1, quercetin-3-glucoside, cycasin, and sorivudine) associated with characterized bacterial enzymes from the original DrugBug publication (Table 1—source data). Both DrugBug and SIMMER were able to predict EC classes for sorivudine, but only SIMMER was able to accurately predict the specific enzyme (BT_4554) responsible for the drug’s degradation. For ginsenoside Rb1 (3.2.1.192), quercetin-3-glucoside (3.2.1.21), and cycasin (3.2.1.21), SIMMER was able to accurately predict EC codes out to sub-sub-class (3.2.1.-), serial designation (3.2.1.21), and sub-sub-class (3.2.1.-) respectively, which was a resolution improvement over DrugBug (Table 1—source data).

We next queried the 28 EC annotated drug-metabolism positive controls against MicrobeFDT (which uses a chemical graph to predict EC codes, but not enzymes) in two ways: first by looking for direct enzyme metabolism events, and second, by looking for enzyme metabolism of compounds that overlap chemically with the positive control in question. When directly queried, MicrobeFDT produced metabolism predictions for four of the 28 positive controls, three of which were correct. When queried with chemically similar compounds according to its graph, MicrobeFDT produced metabolism predictions for 13 of the 28 positive controls, and four were correct (14% overall accuracy in comparison to SIMMER’s 93%). It is important to note that MicrobeFDT is reliant on a fixed database that cannot be modified by the user, meaning that a compound cannot be queried if it is not already present in MicrobeFDT’s graph. We finished the comparison between SIMMER and MicrobeFDT by querying SIMMER with the metabolism use-case described in the MicrobeFDT publication, altretamine demethylation. In our hands, there was no Cypher query against the MicrobeFDT database that resulted in a demethylase EC code (we determined possible demethylase EC codes by running a query in the Swiss Institute for Bioinformatics Enzyme Nomenclature Database)(Bairoch, 2000). We performed queries of direct EC annotation for melamine and altretamine, as well as EC annotation queries for any compound with either substructure or toxicity overlap with altretamine or melamine. The closest result to a demethylase enzyme was a cypher query of toxicity overlap with altretamine that yielded a nitric oxide synthase (EC 1.14.13.39) acting on L-arginine among its results (Table 1—source data). For its significant EC (reaction type) prediction, SIMMER identified altretamine demethylation appropriately as an oxidoreductase reaction acting on a CH-NH group of donors (EC 1.5.-), but not significantly as a demethylation event (Table 1—source data).

This comparison illustrates SIMMER’s enhanced accuracy over other methods for the use case of characterized drug metabolism events by gut bacteria, and also illustrates SIMMER’s novel ability to predict chemical transformations not previously described in literature or databases.

### SIMMER predicts novel drug-metabolizing enzymes

After establishing SIMMER’s accuracy in predicting drug-metabolizing enzymes in the human gut environment, we predicted EC codes, functional annotations, and enzyme sequences for novel microbiome drug metabolism reactions that do not yet possess a responsible, characterized enzyme (Figure 6—source data). From our literature curation of 298 non-antibiotic therapeutics affected by the microbiome (Supplementary File 1), we were confident that 88 are directly metabolized by gut bacteria due to their association with an identified bacterial metabolite in the literature. We formatted these 88 reactions in SMILES format and input them as queries to SIMMER.

Of the 88 reactions queried, SIMMER determined significant EC predictions for 75 reactions (86.2%), and 61 (70.1%) of these were out to the serial designation (i.e. highest resolution) EC code (Figure 6—source data). This list of 61 transformations presents reactions for which we believe enzyme characterization is worth pursuing as our predictions are significantly similar to enzymes already explored in the literature. SIMMER’s EC predictions resulted in expanded and diversified EC class membership for drug-transformations known to occur in the microbiome (Figure 6C). Of interest, this analysis resulted in a large expansion of putative hydrolysis, reduction, and isomerization reactions in the human gut microbiome. The number of SIMMER predictions varies widely by reaction, with median output of 372 genes, 286 genomes, and 10 phyla predicted as responsible across the 88 reactions (Figure 6—source data, Figure 6A). Unsurprisingly, many of these reactions are predicted to occur due to enzymes found in Firmicutes, Bacteroidetes, Actinobacteria, and Proteobacteria, but there are also SIMMER enzyme predictions in phyla not previously associated with drug metabolism (Figure 6B).

**Figure 6.**
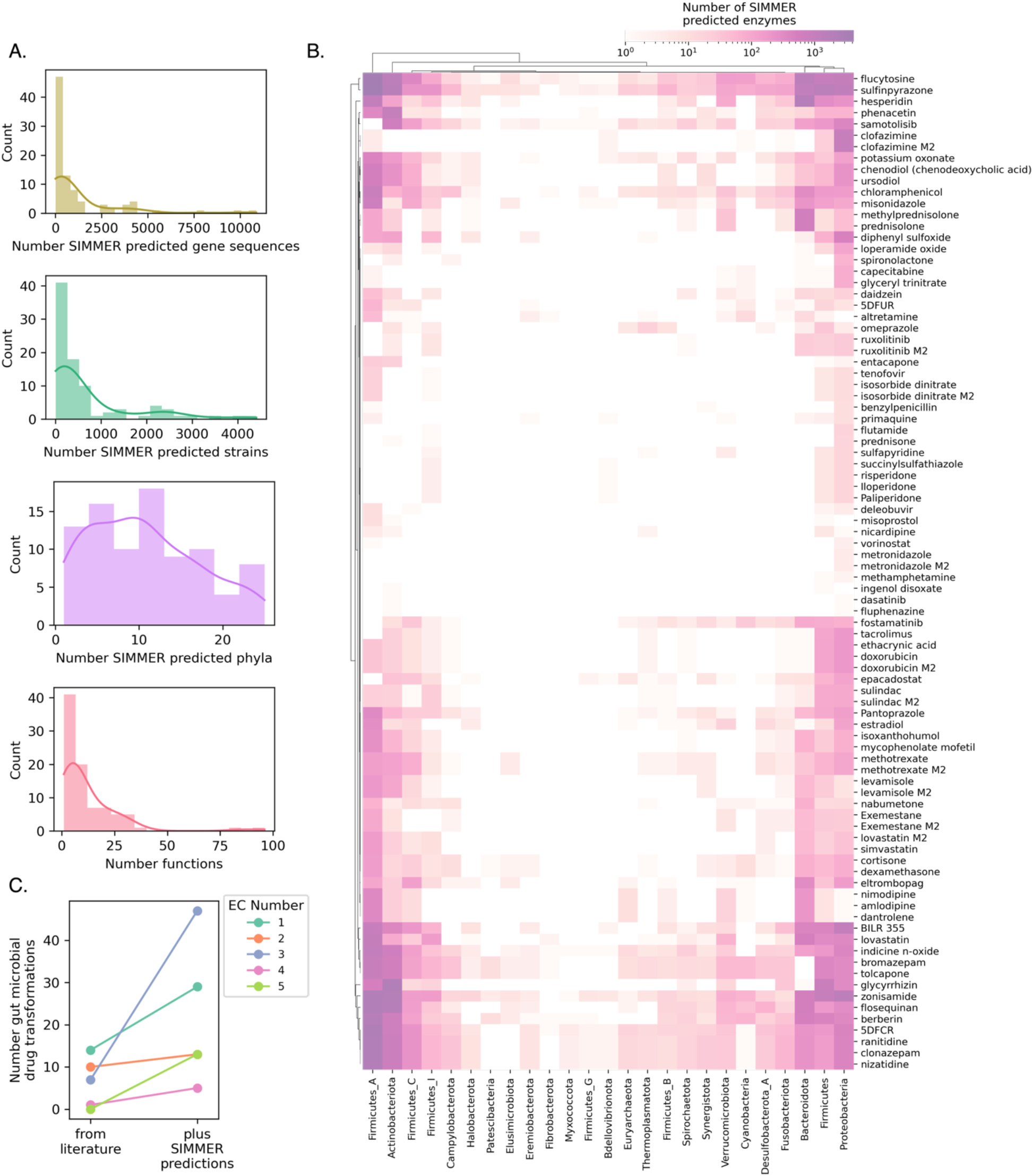
SIMMER predicts novel drug metabolizing enzymes. **(A)** Distributions depict the unique number of genes, strains, and phyla predicted to be responsible for 88 reported drug transformation reactions, as well as predicted gene functions. **(B)** A heatmap illustrating the number of phyla from the UHGG database capable of performing 88 drug metabolism events. Color intensity refers to the number of unique drug-metabolizing enzymes for a given phylum conducting a given reaction. **(C)** Enzyme Commission Class representation for bacterial transformations of therapeutics before and after the employment of SIMMER. Our predictions greatly expand the number of characterized reduction (EC1), hydrolysis (EC2), and isomerization (EC5) events and modestly increase the number of transferase (EC2) and lyase (EC4) events.

Eight of the 88 novel transformations were among those investigated in a high-throughput study exploring the metabolism of 571 compounds in *ex vivo* stool samples (Javdan et al., 2020). This publication demonstrated bacterial degradation of 57 therapeutics in a single pilot donor stool sample (with associated shotgun sequencing), as well as in 20 human stool samples (with associated 16S rRNA gene sequencing). While this study greatly expanded the number of drugs known to break down in the presence of gut bacteria and identified eight metabolite structures, it only identified a responsible enzyme in two of the 57 drug degradation cases due to the low-throughput nature of enzyme characterization. To further assess SIMMER’s ability to predict novel enzymes, and to demonstrate the utility of using SIMMER in an experimental context, we investigated the presence of our predictions in the Javdan, et al. study sequencing results. Because shotgun metagenomics sequencing for the pilot donor was deposited, we were able to confirm via tBLASTn searches that SIMMER enzyme predictions were directly found in the pilot donor stool sample for all eight of the reactions with identified metabolites (Figure 7—source data). However, the sequencing data from the 20 human donor study was only 16S profiling, so we were unable to look directly for SIMMER enzyme predictions. We were able to ensure that genomes found in metabolizing stool samples contain SIMMER predictions. We found that donors who could metabolize a given drug possessed a significant enrichment of genomes that contain enzymes predicted by SIMMER. This was the case for five out of the six reactions analyzed (Figure 7—source data, Figure 7A).

**Figure 7.**
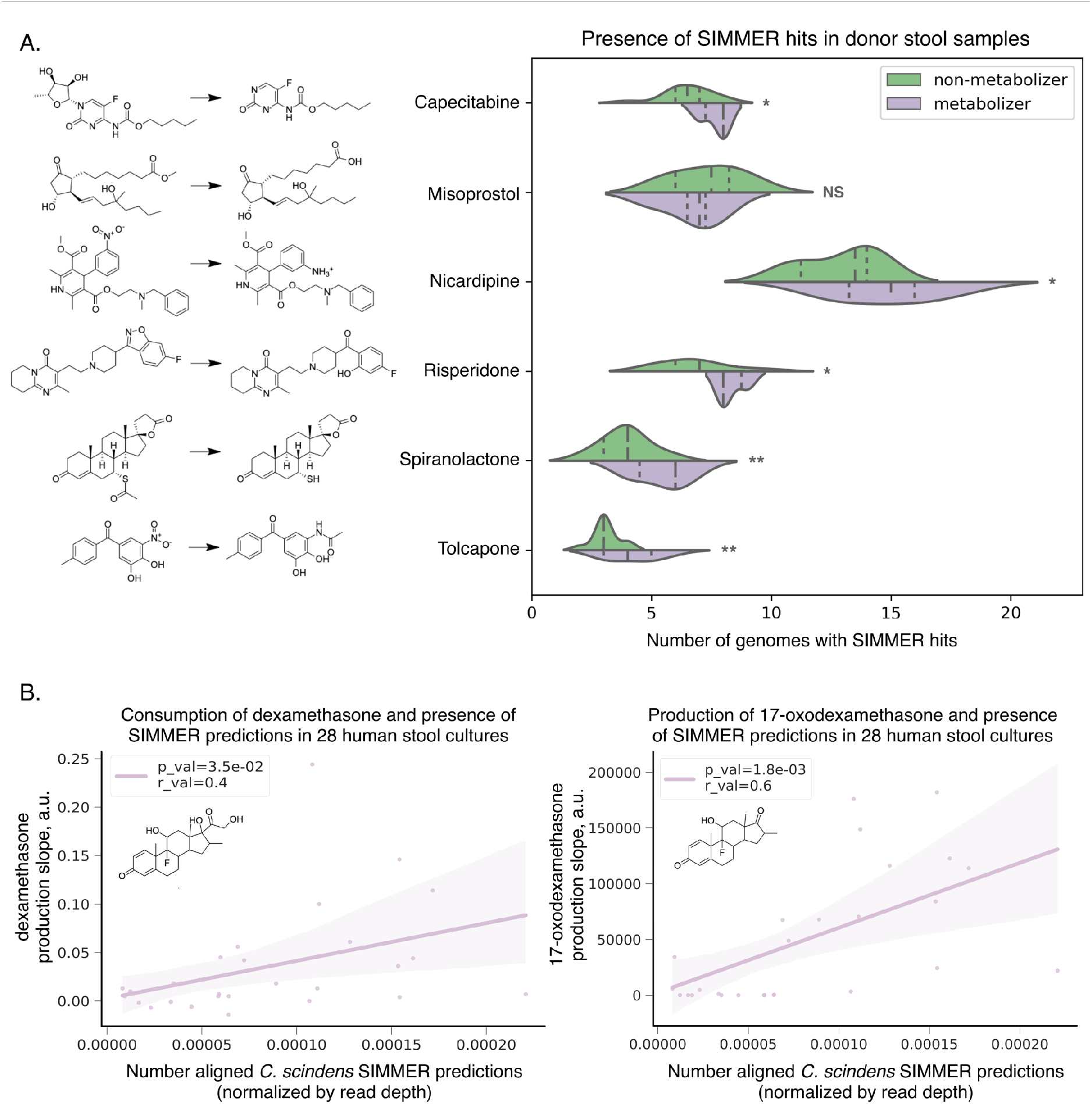
SIMMER predicted enzymes explain inter-individual variations in drug metabolism. **(A)** Donors (N=20) from the Javdan, et al. 16S rRNA gene sequencing study (Javdan et al., 2020) possessed an enrichment of genomes harboring SIMMER enzyme predictions when metabolism of a given drug was observed. Violin plot curves were made using a seaborn package that performs a kernel density estimation of the underlying datapoint distribution. Chemical transformations were drawn using ChemDraw software. Single asterisks denote p-values ≤ 0.05, and double denote p-values ≤ 0.01. **(B)** There was a significant correlation between a human stool sample’s ability to consume dexamethasone (consumption slope, a.u.), to produce 17-oxodexamethasone (production slope, a.u.), and the number of aligned SIMMER predicted sequences for side chain cleavage of dexamethasone. Patient (N=28) conversion slopes and metagenomics data were accessed from the original study (Zimmermann et al., 2019). Chemical structures were drawn using ChemDraw software.

Among the 88 novel transformations was the side-chain cleavage of dexamethasone to 17-oxodexamethasone. Dexamethasone was recently shown to be metabolized solely by *Clostridium scindens* (ATCC 35704) out of a collection of 76 isolates representative of the human gut microbiome (Zimmermann et al., 2019b). When dexamethasone metabolism was assessed in 28 human stool samples, metabolite formation varied substantially by individual, but could not be explained by *C. scindens* species abundance. We sought to understand this lack of correlation. To do so, we assessed the abundance of *C. scindens* SIMMER predictions across the 28 samples (i.e., the amount of enzyme in genomes from different strains, not the amount of the species). We found a significant association between metabolite formation and number of SIMMER enzymes, and also a significant association between parent compound consumption and number of SIMMER enzymes (Figure 7B).

It came to our attention while preparing this manuscript that recombinant steroid-17,20-desmolase (DesAB) enzymes from *C. scindens* were shown to perform side-chain cleavage on prednisone, but also to a lesser extent on dexamethasone. DesAB’s reduced activity for dexamethasone was assumed to be due to the compound’s potentially inhibitory 16α-methyl group (Ly et al., 2020). To ensure that SIMMER’s enzyme prediction for dexamethasone cleavage was not enriched in metabolizing stool samples due to co-occurrence with already known DesAB, we next assessed the abundance of *desAB* reads across the 28 samples, and found no significant correlation between number of reads and either metabolite formation or dexamethasone consumption slopes (Figure 7—figure supplement 1).

These results indicate that species level information alone is not enough to predict chemical transformations in a microbiome sample, but with SIMMER, knowledge of responsible enzymes can recapitulate a sample’s potential for therapeutic degradation.

### SIMMER software

In addition to providing SIMMER (https://github.com/aebustion/SIMMER) as a command line tool that quickly generates enzyme sequence predictions (fasta and tab-separated-value files), EC predictions (tab-separated-value file), and MetaCyc reactions ranked by similarity (tab-separated-value file) based on a user’s input reaction, SIMMER is also available as a user-friendly website (https://simmer.pollard.gladstone.org/). The user can either input one query reaction at a time, or upload multiple reactions in tsv file format (Figure 8). All output types available with the SIMMER command line tool are likewise retrievable via the SIMMER website.

**Figure 8.**
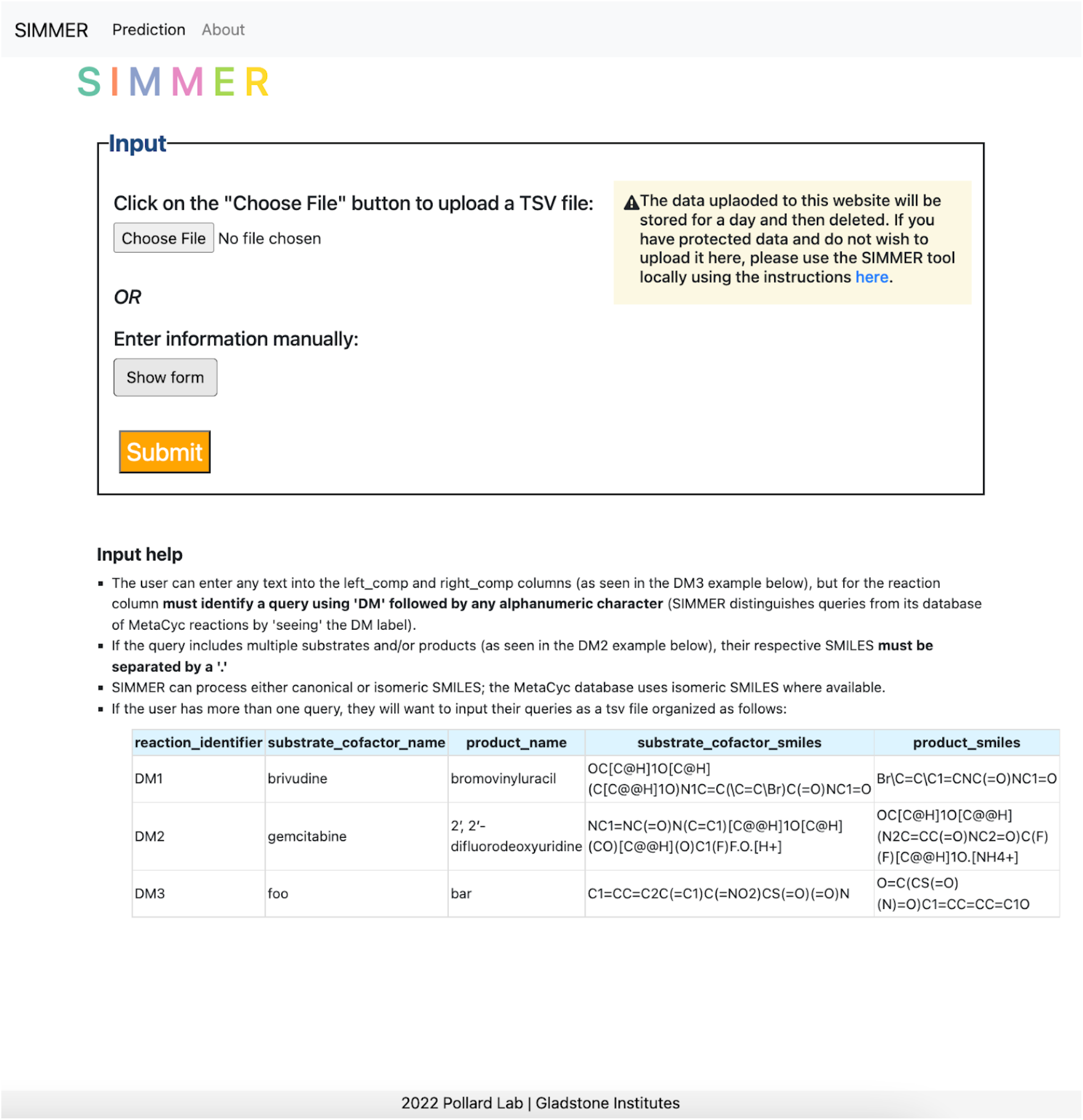
SIMMER webtool. The landing page for the SIMMER website (https://simmer.pollard.gladstone.org/) allows the user to upload a TSV file of queries or add a single query manually to run SIMMER on. It is recommended to use the command-line tool (https://github.com/aebustion/SIMMER) for more than 10 input queries.

## Discussion

In this work, we created a tool that appropriately describes reaction chemistry and harnesses all current information on gut bacterial sequences, both from isolates and metagenome assembled genomes. This advances our ability to discover chemical transformations in the human microbiome, because previous methods for *in silico* metabolism prediction had several key limitations, including low accuracy. Here, we demonstrated SIMMER’s ability to recover known drug-metabolizing enzymes in the human gut, to extend previous experimental findings for multiple drug metabolism events by identifying candidate enzymes, and to add clarity to the genetic component of dexamethasone metabolism by *C. scindens*.

To describe chemical reactions, we were initially influenced by recent research that employed substrate and product chemistry to compare bacterial-drug metabolism events to primary reactions in the MetaCyc database, but without the end-goal of EC and enzyme identity prediction (Mallory et al., 2018). From a reaction description standpoint, the published method was still limited in that it only included a description of one substrate and one product per reaction, precluding it from utilizing cofactors and from accurately describing anything other than intramolecular rearrangements (EC class 5, Figure 2—figure supplement 3). For this reason, we employed a chemical representation technique that can describe multiple inputs and outputs for a single reaction (Schneider et al., 2015).

To connect these chemical descriptions to bacterial proteins in the human gut, we knew it was important not to rely on EC codes (as previous methods have done) to find relevant sequences. While EC codes are helpful for describing reaction types, from an enzyme perspective they contain no information about substrate specificity for a particular compound. We instead chose to create sequence searches of large genome databases directly from enzymes known to conduct chemically similar reactions, whether or not they have been fully annotated with an EC code. For example, enzyme BT_4096 responsible for diltiazem deacetylation (Zimmermann et al., 2019b) is not yet annotated by the EC, yet SIMMER was able to accurately predict the deacetylase enzyme responsible for diltiazem metabolism because it does not require EC annotation for its enzyme predictions. Indeed, instead of relying on EC codes for sequence searches, we harnessed EC annotations in the MetaCyc database to create a novel EC predictor. Using this tool, we accurately predicted diltiazem deacetylation as an EC3 hydrolysis reaction. While EC prediction methods based on sequence exist, to our knowledge, this is the first instance of an EC prediction method based solely on the chemical description of a reaction.

SIMMER achieved high accuracy when applied to known drug-metabolism events in the gut microbiome. Correct EC designations and enzyme sequences were recovered for 31 drug metabolism events previously characterized in microbiome literature. These reactions span multiple EC classes, and were described by multiple publications, demonstrating the wide application and accuracy of SIMMER. While SIMMER provides high accuracy (i.e. true positive) enzyme predictions for chemical transformations in the human gut, the potential for false positives may be high, as its enzyme lists are not filtered by biologically relevant metrics like substrate affinity or flux consistency in a microbial community. To the former point, users may wish to employ tools like Similarity Ensemble Approach to narrow in on hits most likely to interact with compounds of interest (Keiser et al., 2007). To the latter, a user could choose to further analyze their SIMMER output for flux-balance if the predicted SIMMER bacterial species are described in current metabolic reconstruction models (Heinken et al., 2020; Magnúsdóttir et al., 2017).

Due to its high accuracy predictions for previously described drug-metabolism events, we also used SIMMER to predict novel drug metabolism chemistry in the human gut, and expanded what we know of the bacterial enzymes at play in identified drug transformations by gut bacteria. Recent high-throughput experimental research has greatly increased our knowledge of the number of drugs altered by bacteria in the human gut, but has led to a bottleneck in identifying the responsible bacterial enzymes. While direct experimentation is a necessary component to elucidating the bacterial players responsible, *in silico* methods like SIMMER are needed to help prioritize which of the many bacterial species and enzymes to assess. Here we showed that SIMMER both corroborates previous high throughput experimental data, and also adds increased clarity to the findings. While a previous experimental study was able to elucidate the importance of an isolate *C. scindens* in the metabolism of dexamethasone, the abundance of *C. scindens* in human samples did not correlate with metabolism. When assessed with SIMMER, however, a significant correlation between metabolite production and amount of SIMMER predictions was observed. This finding demonstrates that species identity alone is not enough to explain bacterial chemical transformation, and that responsible genetic elements must be interrogated as well.

Two previous computational tools exist for describing non-antibiotic microbial drug metabolism. MicrobeFDT groups thousands of compounds based on their similarity to one another and annotates compound groups based on any known links to EC numbers, and subsequently, microbes known to contain such EC codes (Guthrie et al., 2019). This network approach was an important addition to the exploration of microbiome metabolism, but its use is limited to a fixed database of chemicals and EC annotations which prevents the user from exploring novel chemistry and also from utilizing hypothetical protein data gathered from metagenomic sequencing studies. Furthermore, MicrobeFDT’s accuracy within its database of substrates is limited by its exclusive description of substrates rather than full reactions. DrugBug, a tool that employs Random Forests rather than a network approach, also exhibits limited power and accuracy due to its sole reliance on substrate chemistry and relatively small database of only 491 isolated bacterial genomes from the human gut (Sharma et al., 2017). Of note, our comparison of SIMMER’s performance to existing methods necessitated downloading and analyzing our positive control list against the other tools, as none of the previous publications provided any computational validation or accuracy metrics.

One user pitfall of SIMMER in comparison to previous methods, is that a reaction’s product(s) and cofactor(s) identity is required to achieve the high accuracy enzyme prediction described here. This is a limitation, as a growing amount of LC-MS/MS data in microbiome research only reports whether or not a compound is depleted in the microbiome and the mass/charge ratio of the product formed, not the product identity. While it is technically possible for the user to submit a SIMMER query that only consists of a substrate, or uses a compound identity as both substrate and product, we do not recommend this due to the previously discussed lack of accuracy when only considering substrates (Figure 2—figure supplement 3, Table 1—source data). For users wishing to utilize SIMMER with a compound of interest and its either unknown or uncharacterized products, additional tools such as BioTransformer could be used in tandem to create product template predictions before querying (Djoumbou-Feunang et al., 2019). Lastly, if the user does not hold a certain level of knowledge in chemistry, appropriate cofactors (such as water employed in a hydrolysis event) might be omitted from a query, leading to lower accuracy predictions. If a user is unsure which cofactors may be at play in their reaction of interest, reaction rules tools such as RetroRules could be employed (Duigou et al., 2019).

Another limitation of SIMMER is that its underlying protein data is solely metagenomics data from the human gastrointestinal tract, but some compounds, such as the vaginal gel tenofovir, are known to be altered by bacteria in non-GI tract settings (Klatt et al., 2017). That being said, for transformations in the human gut, SIMMER employs the largest available database of relevant bacterial sequences, and the tool could easily be expanded in the future to include other human body sites as well as non-host associated environments. Further related to database constraints, while SIMMER is novel in its ability to query reactions not previously described in chemistry databases, its search space is still limited to reactions that broadly relate to those captured in MetaCyc. As MetaCyc expands, or additional databases get employed, SIMMER will likely be able to make increasingly fine-tuned predictions.

SIMMER enters microbiome biotransformation research at an important point: while there are hundreds of microbiome altered compounds which are in need of enzyme identification, there are also a sufficient number with characterized enzymes to enable us to test the tool’s accuracy. Its ability to predict these known enzymes accurately builds confidence for its predictions of yet unknown enzymes. With this tool in hand, microbiome researchers can make informed hypotheses before embarking on the lengthy laboratory experiments required to characterize novel bacterial enzymes that can alter human ingested compounds. Continued refinement of SIMMER and other computational tools will accelerate microbiome research, providing data-driven hypotheses for experimental testing and a first step towards understanding the full scope of metabolism by the human microbiome.

## Materials and methods

### Preparation of SIMMER’s underlying chemical data

13,387 gene annotated bacterial reactions were downloaded from MetaCyc (Caspi et al., 2008). Each reaction from the database contained Simplified Molecular Input Line Entry System strings (SMILES) of reactant(s) and product(s), EC code if available, and UniProt or Entrez identifiers for sequences that catalyze the reaction. All MetaCyc compounds were then protonated based on the pH environment of 7.4 in the human small intestine, where most oral drug absorption occurs. Protonation states were calculated using ChemAxon’s cxcalc majorms software (“cxcalc calculator functions,” n.d.).

RDKit’s rdChemReactions module was employed to create chemical fingerprints representing each MetaCyc reaction. Chemical reaction objects were constructed from reaction SMILES arbitrary target specification (SMARTS) strings. Fingerprints for these reactions were then created using the resulting difference of product(s) and reactant(s) Atom-Pair fingerprints (Schneider et al., 2015). SIMMER users can also opt to use Topological Torsion, Pattern, or RDKit fingerprints, but unless otherwise stated, all analyses in this manuscript use Atom-Pair difference fingerprints. Of the 13,387 MetaCyc reactions, 8,914 were able to be fingerprinted using this method. Failed fingerprints were due to ambiguous SMILES identifiers or presence of non-small molecule compounds in a reaction, such as peptides.

After creating fingerprint vectors for all MetaCyc reactions, an 8,914 by 8,914 pairwise similarity matrix of Tanimoto coefficients was created. These Tanimoto vectors make up SIMMER’s underlying chemical data.

### Preparation of SIMMER’s underlying protein data

For each of the 8,914 fingerprinted MetaCyc reactions, all relevant gene sequences were retrieved from the MetaCyc reaction’s UniProt and Entrez database linkouts. If at least two genes, with a median pairwise sequence similarity greater than or equal to 27%, were linked to a given MetaCyc reaction, the sequences were used to create a multiple sequence alignment and subsequent profile hidden Markov model (pHMM) using Clustal Omega and HMMER3 (version 3.2.1) software respectively (Eddy, 2009; Sievers and Higgins, 2014). This similarity cutoff was chosen based on previous protein family literature (Mi et al., 2021). If fewer than two genes, or genes with less than 27% global similarity, were associated with a given MetaCyc reaction, a pHMM of the MetaCyc gene(s) PANTHER subfamily was retrieved via InterPro linkouts (Mi et al., 2021). MetaCyc derived and PANTHER subfamily pHMMS were then queried against a Unified Human Gastrointestinal Genome (UHGG) collection of 286,997 isolate genomes and metagenome assembled genomes from the human gut environment using the HMMER3 hmmsearch module (Almeida et al., 2020; Eddy, 2009). In the case of MetaCyc reactions with too few sequences, too low a median pairwise sequence identity, *and* a missing PANTHER database subfamily pHMM, single sequence protein queries were conducted against the UHGG databse using HMMER3’s phmmer module, which internally created protein profiles for the single query sequences based on a position-independent scoring system. Resulting enzyme hit lists were filtered to only include high significance hits (e-value < 1E−5, and hit length >= half of the input pHMM alignment or single sequence length). In sum, for each MetaCyc reaction, a profile representing the diversity of the enzyme family for that chemical transformation was used to find sequence similar hits in the human gut microbiome that can mediate chemically similar reactions.

Each human gut microbiome hit was further described by the identity, prevalence, and abundance of the bacterial strain in which it resides. To establish prevalence and abundance of UHGG strains, metagenomic analysis was performed on the Predict (Personalised Responses to Dietary Composition Trial) cohort due to its high number of samples and favorable sequencing depth (Asnicar et al., 2021). Shotgun metagenomic reads were analyzed with MIDAS2 an implementation of Metagenomic Intra-Species Diversity Analysis Subcommands (MIDAS) (Nayfach et al., 2016; Zhao et al., 2022) designed for the UHGG database. Presence of a SIMMER predicted species in a given sample was established when reads mapped (HS-BLASTN) to 15 single-copy universal genes for that species (Chen et al., 2015), with at least 75% alignment coverage. To assess the gene content of a sample, shotgun metagenomic reads were aligned to a MIDAS2 created pangenome of the SIMMER species’ genes clustered at 99% nucleotide identity. Copy number of a SIMMER gene prediction was established by dividing aligned prediction reads by the full length of the prediction. This number was then normalized by the read coverage of 15 single-copy universal genes in the same sample to estimate copy number per cell. Presence of a SIMMER enzyme was established if at least 0.35 gene copies per cell were present in a sample.

Phylogenetic trees were also constructed for each hmmsearch and phmmer result. For each set of MetaCyc reaction human gut microbiome enzyme hits, CD-HIT was used to cluster results at 95% identity (Fu et al., 2012). Then MUSCLE was used to create a multiple sequence alignment for input to FastTree (Edgar, 2004; Price et al., 2009). Compact tree visualizations were made in R using ggtree and ggtreeExtra (Xu et al., 2021; Yu et al., 2017). All tree tips were colored by phylum, and surrounded by circle annotators describing a given hit’s Prokka predicted function, genome type (i.e. from an isolate or metagenome assembled genome), and prevalence/abundance in the Predict cohort (Seemann, 2014).

### Query functionality of SIMMER

The query functionality of SIMMER was designed similarly to the precomputed chemistry data. After receiving an input chemical transformation (or tsv describing multiple input reactions) in the form of SMILES, SIMMER fingerprints the reaction(s) and compares it to the precomputed chemical space by computing the Tanimoto coefficients between the input(s) and all precomputed reactions. The 8,914 precomputed MetaCyc reaction Tanimoto vectors are then sorted by ascending euclidean distance to the query Tanimoto vector. SIMMER by default ranks reactions’ euclidean distances based directly on the Tanimoto vectors, but if a user’s inputs require a decrease in computational burden, PCA can be employed after similarity matrix creation and before euclidean distance rankings. The number of PCs to be used depends on the fingerprint style employed, and was determined by the Kaiser criterion. Unless otherwise stated, all analyses in this manuscript employed the full Tanimoto similarity matrix with no PCA reduction. Human gut microbiome enzymes that may conduct the input reaction are reported from the precomputed UHGG hmmsearch or phmmer results of the closest euclidean distance MetaCyc reaction. Significantly enriched EC identities (i.e. reaction types) are also reported.

### Reaction type predictions

SIMMER predicts an EC code (i.e. reaction type) for a query reaction if there is an enrichment of a particular EC at the top of the reaction list. Enrichment was determined in a manner similar to gene set enrichment analysis (GSEA).(Subramanian et al., 2005) For each EC code associated with MetaCyc reactions, an enrichment score (ES) was calculated by walking down the ranked list of reactions. Starting with a score of zero, each time the given EC is encountered the score increases by one, and each time a different EC is encountered the score decreases by one. At the end of this process, each EC receives an ES that is the score’s maximum distance from zero after walking through the list (Figure 2—figure supplement 1A). Because the MetaCyc database of reactions is unbalanced in its EC code representation, ES scores for a given EC type are divided by the number of times the EC in question occurs in the database. This yields a normalized ES (NES) for SIMMER reporting. Significance is established by comparing the true NES to the NES achieved from 1000 permutations of a shuffled reaction list (Figure 2—figure supplements 1B-C). When multiple EC codes are predicted as significant, they are ranked in ascending order of where in the list of 8,914 reactions the NES occurs. This method was verified by subsampling the database of MetaCyc reactions to equal numbers (N=96) of reactions for each EC class, the broadest resolution level of an EC code. Each of these subsampled reactions was then queried with SIMMER against the entire MetaCyc reaction database to create sorted reaction lists for each query. SIMMER predicted an EC code(s) for each reaction based on the most highly enriched EC. SIMMER’s recall, precision, and accuracy are high for EC class, sub-class, and sub-sub-class level resolution (Figure 2B, Figure 2—source data). For the serial designation of an EC code (the most granular description of an EC code), however, SIMMER’s performance diminished, potentially because enrichment calculations suffer from increased uniqueness in the ranked list and therefore reduced power to determine a match (Figure 2—source data). This indeed appears to be the case; when the database is subsampled to ensure at least three of each unique serial designation, F1-scores (the harmonic mean of precision and recall) and accuracy remain high despite the increased EC resolution (Figure 2—source data, Figure 2—figure supplement 1D).

### Euclidean distance silhouette scores

To analyze SIMMER’s resilience to different reaction chemistry representations, we created a silhouette-like euclidean distance score. For the precomputed MetaCyc chemical dataset of 8,914 reactions (i.e. the Tanimoto pairwise similarity matrix), we split all reactions into their top-level EC codes (i.e. EC class) and determined for each reaction its euclidean distance to all reactions within its EC class versus outside its EC class. From the two distributions (within EC and without EC distances) created, we computed a Kolmogorov-Statistic to determine if the distributions significantly (p<0.05) differed. We repeated this process for finer resolution EC classifications (sub-class, sub-sub-class, and serial designation). Euclidean distance silhouette scores were used to compare different chemical representations, such as fingerprint style, inclusion of products, and inclusion of cofactors.

### Relationship between SIMMER’s underlying chemical and protein data

For MetaCyc enzymes (N=34,279) associated with multiple reactions, one reaction was used as a SIMMER query, and the other reaction(s) searched for in the ordered reaction list output. As a negative control, these reaction similarity results were then compared to all pairwise combinations of MetaCyc enzymes (subsampled to N=34,279) that do not conduct the same reaction.

We also assessed the relationship between chemistry and protein similarity for all pairwise combinations of a subset of MetaCyc reactions annotated with only one protein sequence (N=604 reactions). Chemical similarity was based on the Euclidean distance between two reaction fingerprint vectors in SIMMER’s precomputed chemical space (Figure 1A). Global protein similarity was determined via the Needleman–Wunsch algorithm. The relationship between chemical similarity and protein similarity was assessed with a Pearson’s correlation coefficient and *P* value calculated using a Wald Test with t-distribution of the test statistic.

### Creating a compendium of drug-metabolism use cases from the human gut

To analyze SIMMER under the use-case of drug metabolism, we created a compendium of drug degradations that occur in the human gut microbiome. The compendium of reactions is based on a literature curation of hundreds of papers, and is organized by reactions producing known/unknown metabolites and driven by known/unknown bacterial enzymes. The drug-metabolism positive controls used to assess SIMMER’s accuracy were drawn from the list of reactions possessing a structurally elucidated metabolite and driven by a characterized bacterial enzyme.

We further expanded the positive control list to include sequence-similar enzymes that likely perform the same function. For this expansion, we performed pHMM searches (when a positive control reaction had been characterized with multiple sequence-similar enzymes) and phmmer searches (when a positive control reaction had been characterized with only one sequence) of the UHGG database using HMMER3 software (Almeida et al., 2020; Eddy, 2009). High significance (e-value < 1E−5) hits were kept when the resulting alignment was at least 50% of the input pHMM or sequence length. This list of significant hits was filtered by presence in human ileum or jejunum (the site of human drug absorption) via DIAMOND searches against metagenomic reads from a published study that employed jejunum and ileum endoscopy (Buchfink et al., 2021; Zmora et al., 2018). The hits were also filtered for presence in RNAsequencing data via DIAMOND searches of rnaSPAdes assembled reads from HMP2 metatranscriptomics control patient samples (Bushmanova et al., 2019; Integrative HMP (iHMP) Research Network Consortium, 2019). The hits were lastly filtered by predicted affinity for their substrates using the Similarity Ensemble Approach (Keiser et al., 2007).

### Corroboration of previous high-throughput experimental findings

For the first experimental validation of SIMMER, we analyzed results from sequencing studies (NCBI BioProject: PRJNA593062) described in a previously published high-throughput investigation of bacterial drug metabolism in human stool samples (Javdan et al., 2020). The first sequencing set in this publication was a deep metagenomic sequencing of one pilot individual’s *ex vivo* stool originally evaluated for its ability to degrade hundreds of therapeutics. We used MetaSPAdes with default settings to assemble the metagenomics reads into scaffolds (Nurk et al., 2017). We then queried SIMMER with eight reactions that were structurally elucidated (via nuclear magnetic resonance) by the previous publication, and ensured via TBLASTN searches that SIMMER predicted hits were found in the assembled metagenomic reads. The second sequencing set was a 16S rRNA sequencing experiment of twenty human donor stool samples originally evaluated for their inter-individual variation in bacterial drug degradation. We queried SIMMER with five of these reactions possessing structurally elucidated metabolites, and evaluated enrichment of SIMMER predicted bacterial species in metabolizing versus non-metabolizing donors. Species matches between SIMMER predictions and the 16S study were made using the SequenceMatcher class from the difflib python module set to an 80% ratio cutoff. Enrichment of SIMMER predicted bacterial genomes was then assessed by computing a t-test for number of SIMMER genomes in metabolizers versus number of SIMMER genomes in non-metabolizers for a given reaction.

For experimental corroboration of dexamethasone metabolism, we accessed shotgun sequencing data (PRJEB31790) from a cohort of 28 human stool samples shown to metabolize dexamethasone to varying degrees (Zimmermann et al., 2019b). Shotgun reads were assembled using MetaSpades with default settings. Presence of SIMMER enzyme predictions was established via search with DIAMOND and normalized by sample read depth. Significance was established with a Pearson’s correlation coefficient and *P* value calculated using a student’s *t*-distribution.

### Web tool creation

We used the python web framework Flask (https://flask.palletsprojects.com/en/2.1.x/) to make SIMMER available as a user-friendly website. The website accepts either a single query reaction or multiple query reactions via a file upload and provides the same outputs as the SIMMER command-line tool. The website also allows the user to download the outputs of interest. Keeping in mind the privacy and security of the data that a user might upload to the website, the website is designed to delete all uploaded data within 24 hours from the server. This will ensure security of the uploaded data.

## Supporting information

Supplementary File 1 - literature curation

Supplementary File 2 - SMILES used as SIMMER input

## Data availability

Data generated and analyzed during this study are provided in Figures 2-7 source data files, Table 1 source data file, supplemental files, and at https://github.com/aebustion/SIMMER. Accession numbers of previously published datasets are provided in the Materials and Methods section. SIMMER code can either be run at the SIMMER website (https://simmer.pollard.gladstone.org/) or downloaded directly from the above-linked GitHub.

## Acknowledgements

We thank Russ Altman and Emily Mallory for their work that first inspired the comparison of MetaCyc reactions to known gut microbiome chemistry (Mallory et al., 2018), and Russ Altman for further insightful conversation. We thank Peter Karp and Peter Midford of MetaCyc for providing us with an expanded set of links between MetaCyc reactions and Uniprot or Entrez sequences. We thank Ben Guthrie and Renuka Nayak of the Turnbaugh lab for collaboration and guidance in the ongoing experimental validation of SIMMER enzyme predictions. We thank Daniela Arce and Xiaofan Jin of the Pollard lab for guidance on assessing enzyme presence in the jejunum and ileum of the human intestinal tract. We thank Chunyu Zhao and Jason Shi of the Pollard lab for guidance on the appropriate use of MIDAS2 and the UHGG database. We thank Patrick Bradley for guidance on the use of HMMER3 software in a microbiome drug-metabolism context. We thank members of the Pollard and Turnbaugh laboratories for their suggestions during the research process and writing of this manuscript. Annamarie Bustion was supported by a trainee pilot award from the UCSF Benioff Center for Microbiome Medicine, a predoctoral fellowship in informatics from the PhRMA foundation, an Achievement Rewards for College Scientists (ARCS) Scholarship from the ARCS foundation, and Training Grants from the NIGMS of NIH (5T32GM007175-42 and 2T32GM007175-41). This work was also supported by Gladstone Institutes. P.J.T. was supported by the National Institutes of Health (R01HL122593). P.J.T. and K.P. are Chan Zuckerberg Biohub Investigators.

## Supplementary data

**Supplementary File 1** (attached). Literature curated list of drug-metabolism events in the human gut microbiome

**Supplementary File 2** (attached). SIMMER EC predictions for all MetaCyc reactions

## Supplemental figures

**Figure 2—figure supplement 1.**
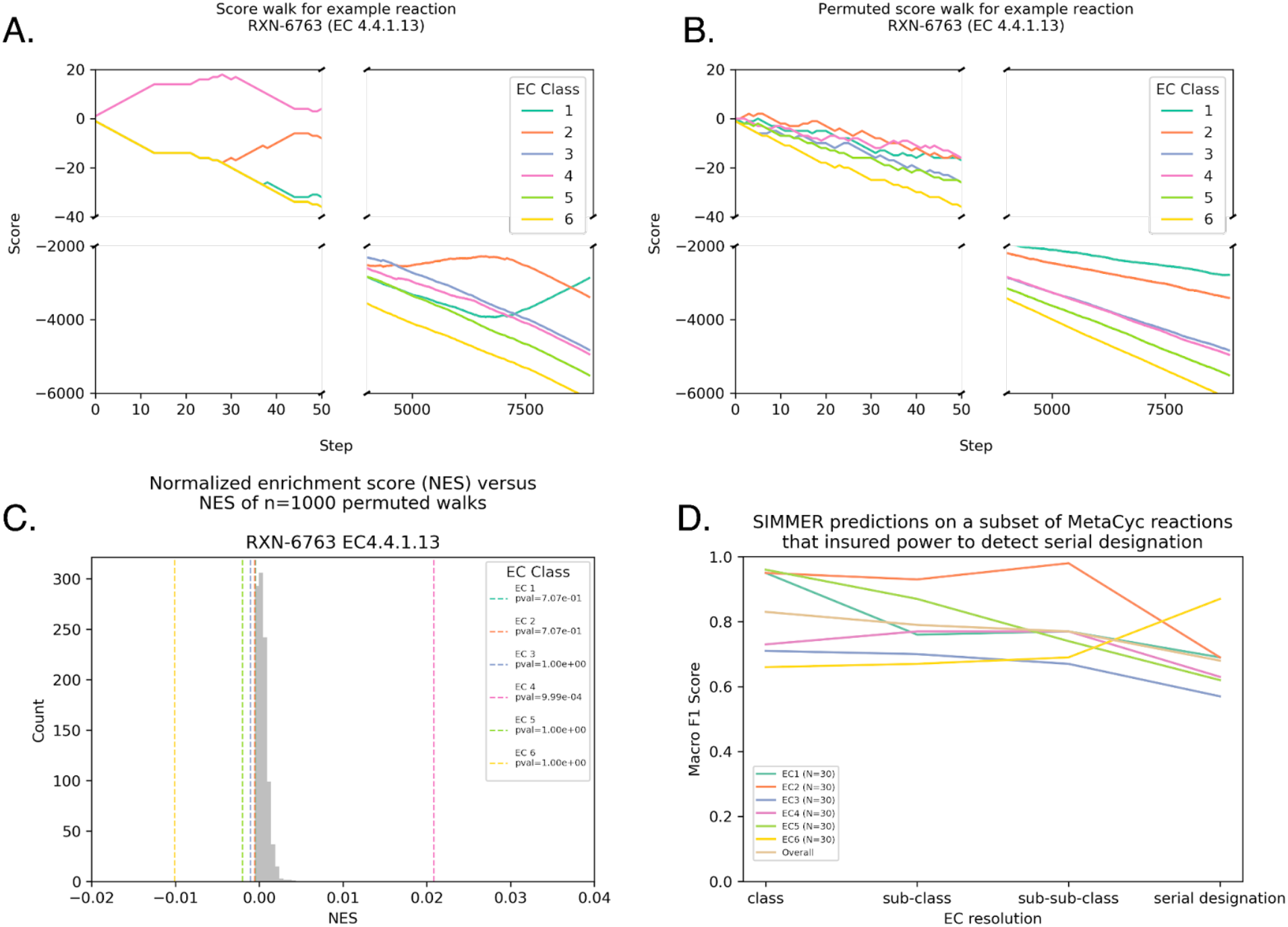
SIMMER predicts an EC code (i.e. reaction type) for a query reaction if there is an enrichment of a particular EC at the top of the reaction list. **(A)** For each EC code associated with MetaCyc reactions, an enrichment score (ES) was calculated by walking down the ranked list of reactions. Starting with a score of zero, each time the given EC is encountered the score increases by one, and each time a different EC is encountered the score decreases by one. Panel A is an example of such a walk with MetaCyc reaction RXN-6763. **(B)** Significance is established by comparing the true ES to the ES achieved from 1000 permutations of a shuffled reaction list. Panel B is an example of how RXN-6763 loses its EC enrichment structure after shuffling. **(C)** Because the MetaCyc database of reactions is unbalanced in its EC code representation, ES scores for a given EC type are divided by the number of times the EC in question occurs in the database. This yields a normalized ES (NES) for SIMMER reporting. This is also performed for the shuffled distributions. **(D)** For the serial designation of an EC code (the most granular description of an EC code), SIMMER’s performance diminished (Figure 2B), because enrichment calculations suffer from increased uniqueness in the ranked list and therefore reduced power to determine a match. This is proven here, as when the database is subsampled to ensure at least three of each unique serial designation, F1-scores (the harmonic mean of precision and recall) and accuracy remain high despite the increased EC resolution.

**Figure 2—figure supplement 2.**
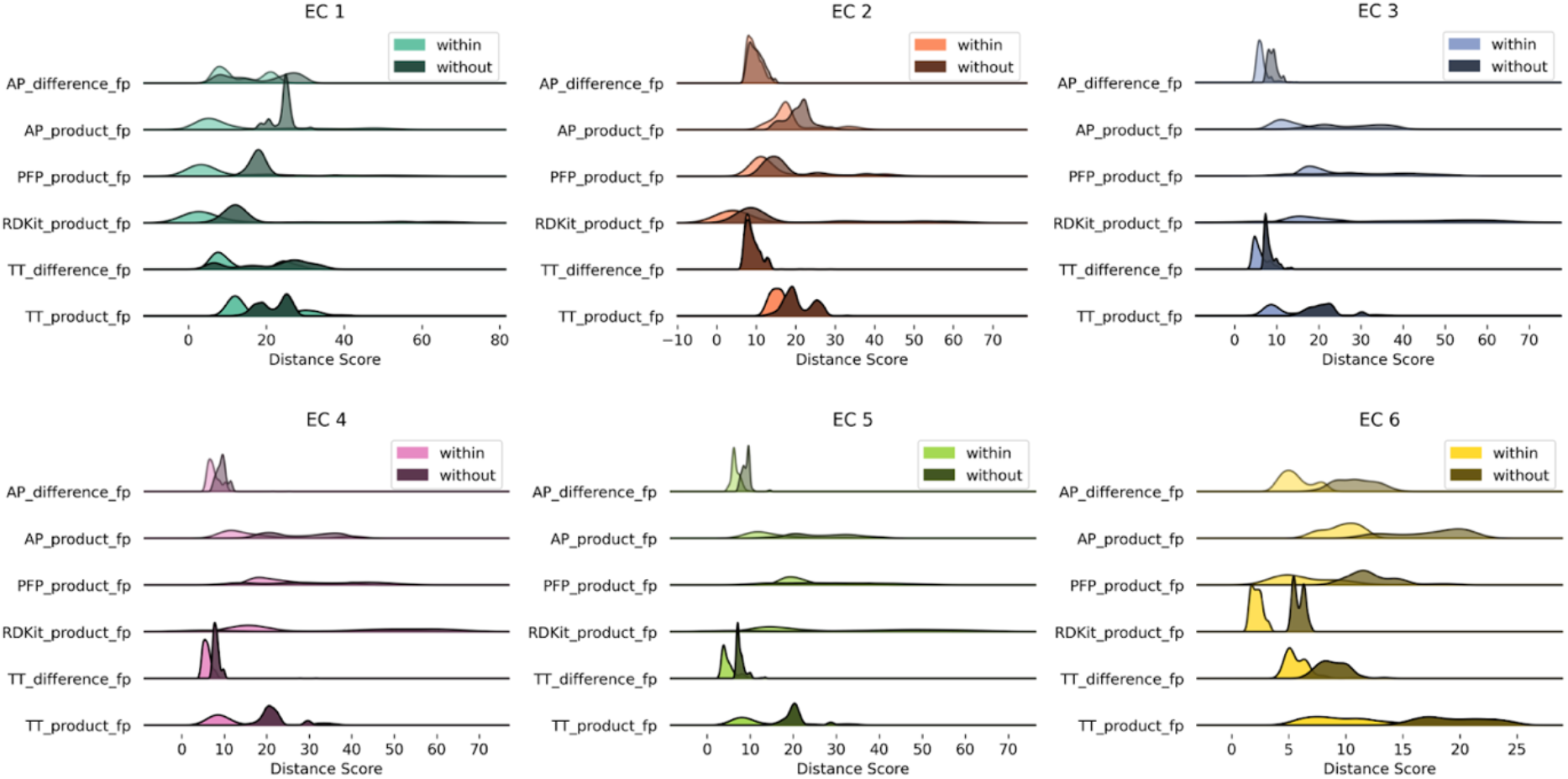
Euclidean distance distributions and silhouette scores for top-level EC codes are resilient to fingerprint type. Distributions are computed as described in Figure 2A.

**Figure 2—figure supplement 3.**
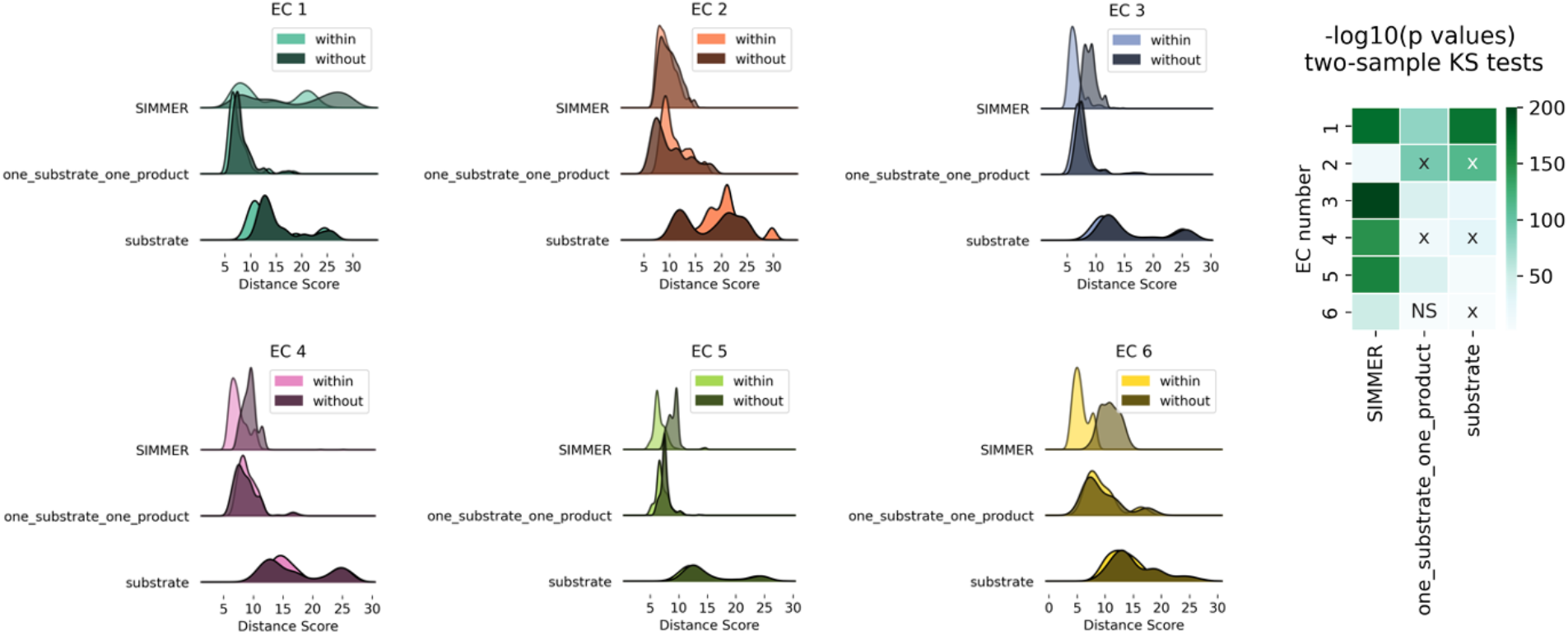
Euclidean distance distributions and silhouette scores for top-level EC codes are sensitive to chemical representation type. Distributions are computed as described in Figure 2A. The resulting score distributions were used to compare SIMMER’s resilience to different chemical representations, such as removal of products and cofactors (the representation methods employed by DrugBug (Sharma et al., 2017) and MicrobeFDT (Guthrie et al., 2019) or the inclusion of exactly one substrate and exactly one product (the Mallory, et al. method (Mallory et al., 2018). For all EC classes, scores were smaller within versus between EC classes using SIMMER’s chemical representation, indicating that SIMMER can detect reaction similarity within EC classes. However, this was not consistently true when using only the input compound without cofactors or products (substrate), or when using only one substrate and product (one_substrate_one_product). From the pairs of distributions, we computed a Kolmogorov-Statistic to determine if the distributions significantly (p<0.05) differed. This showed that the differences were statistically significant for SIMMER chemical representations and reduced, or in the wrong direction, when using a reduced representation. X’s in the KS test heatmap indicate an incorrect direction of difference (without grouping closer than within).

**Figure 4—figure supplement 1.**
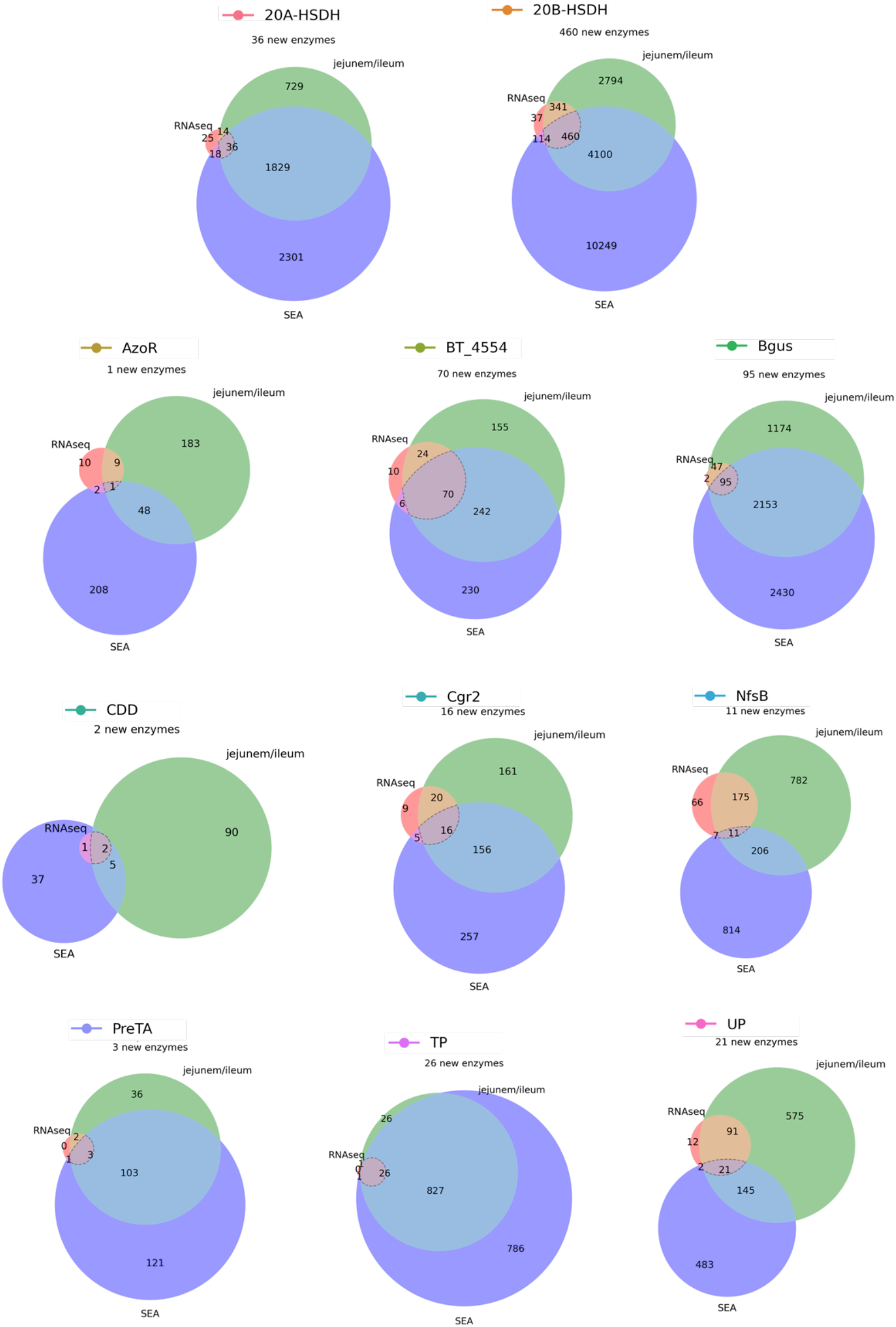
An expanded list of gut bacterial enzymes relevant to known cases of drug metabolism. 52,849 total candidate homologs (a median of 1,087 candidates per enzyme) were obtained via pHMM searches with sequences from the literature. These putative homologs were filtered for presence in the human jejunum/ileum, presence in RNA-sequencing studies, and predicted affinity for the substrate in question using SEA. This resulted in 741 additional enzyme sequences for 31 positive control reactions.

**Figure 5—figure supplement 1.**
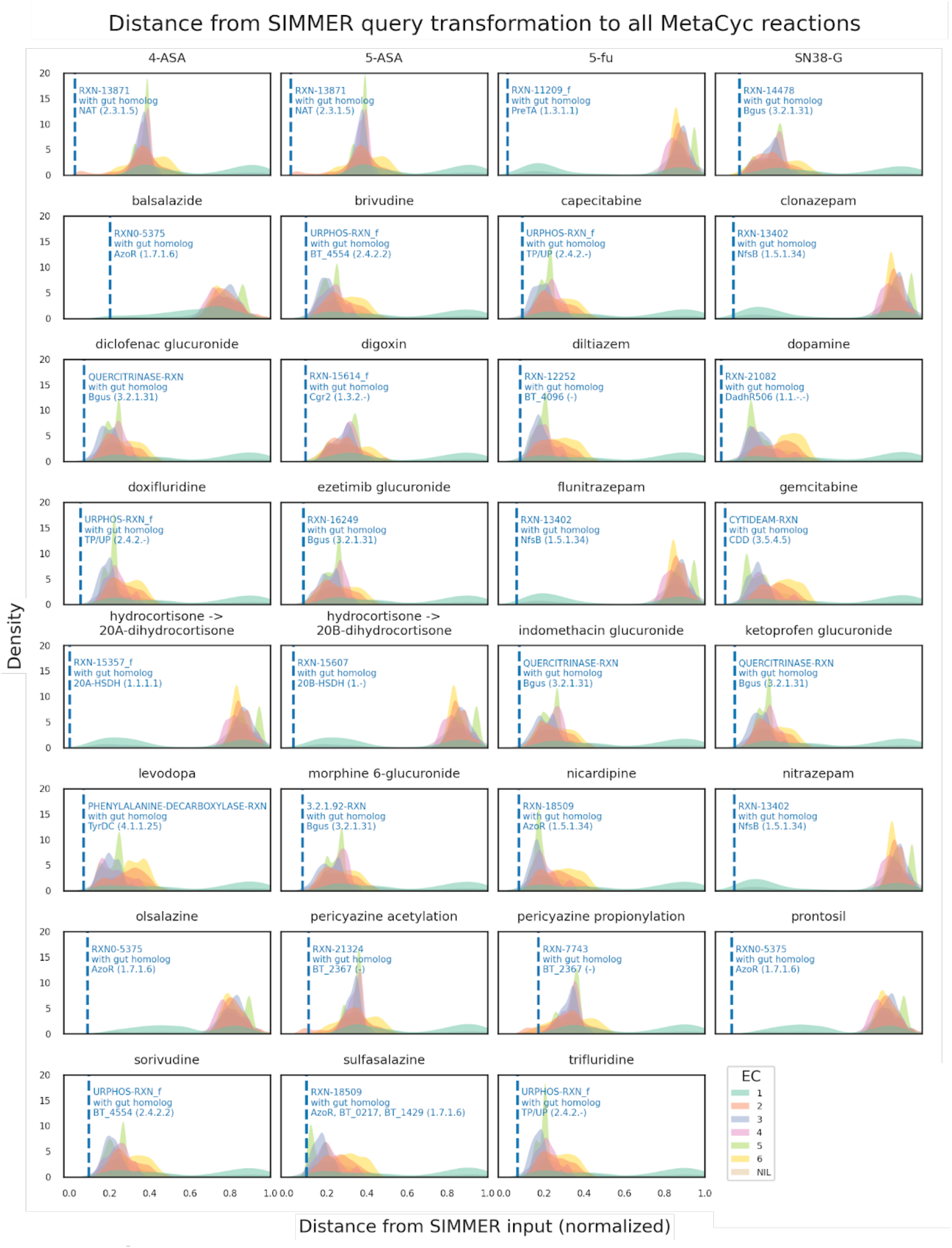
Distributions of all MetaCyc reactions’ euclidean distances to the positive control list queries. For this analysis, each of the 31 positive control reactions was queried to SIMMER. Distributions were created based on all (N=8,914) MetaCyc reactions’ distance to a given input metabolism event, and colored by their EC class annotation in MetaCyc (oxidoreductases, transferases, etc.). The dashed blue line depicts the MetaCyc reaction that yielded a gut microbiome homolog matching the known positive control sequence. For example, when queried with 5-ASA acetylation, SIMMER outputs a most-similar MetaCyc reaction (RXN-13871) that SIMMER linked (via sequence similarity) to an N-acetyltransferase known from previous literature to drive 5-ASA metabolism in the human gut.

**Figure 7—figure supplement 1.**
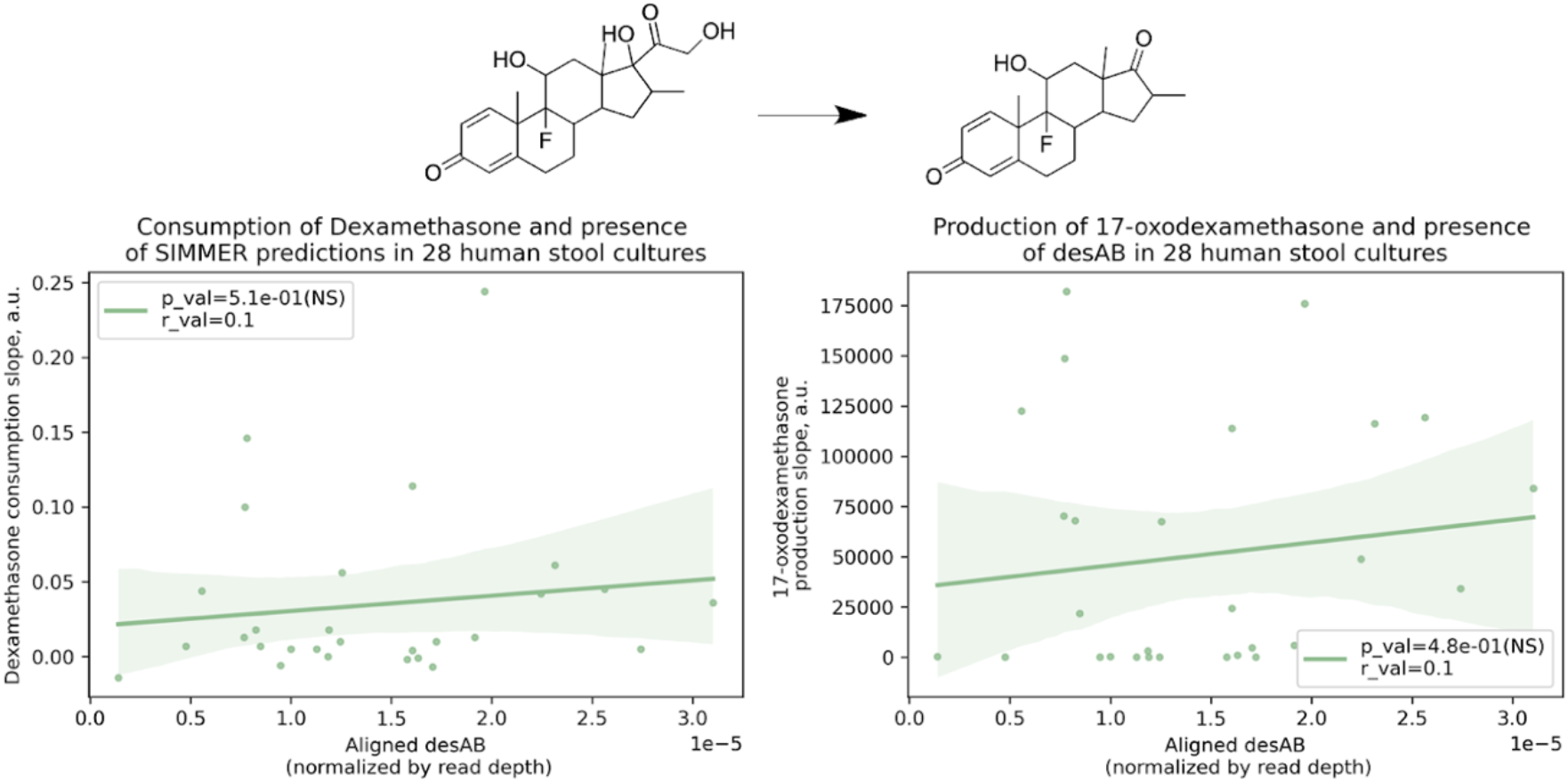
There was not a significant correlation between a human stool sample’s ability to consume dexamethasone (consumption slope, a.u.) or to produce 17-oxodexamethasone (production slope, a.u.), and the number of aligned *desAB* sequences. Patient (N=28) conversion slopes and metagenomics data were accessed from the original study (Zimmermann et al., 2019). This adds confidence to the finding described in Figure 7B, as it means SIMMER predictions were not correlated with dexamethasone metabolism due to co-occurrence in *C. scindens* with a gene previously reported to underlie bacterial side-chain cleavage of steroids (Ly et al., 2020).

